# Towards a neuroscientific understanding of play: A neuropsychological coding framework for analysing infant-adult play patterns

**DOI:** 10.1101/202648

**Authors:** Dave Neale, Kaili Clackson, Stanimira Georgieva, Hatice Dedetas, Sam Wass, Victoria Leong

**Affiliations:** University of Cambridge, UK; University of East London, UK; Nanyang Technological University, Singapore

**Keywords:** Play, mother-infant interaction, neuroscience, coding, social interactions

## Abstract

During early life, play is a ubiquitous activity, and an individual’s propensity for play is positively related to cognitive development and emotional well-being. Play behaviour is diverse and multi-faceted. A challenge for current research is to converge on a common definition and measurement system for play ‒ whether examined at a behavioural, cognitive or neurological level. Combining these different approaches in a multi-level analysis could yield significant advances in understanding the neurocognitive mechanisms of play, and provide the basis for developing biologically-grounded play models. However, there is currently no integrated framework for conducting a multi-level analysis of play that spans brain, cognition and behaviour. The proposed neuropsychological coding framework uses grounded and observable behaviours along three neuropsychological dimensions (sensorimotor, cognitive and socio-emotional), to compute inferences about playful behaviour and related social interactional states. Here, we illustrate the sensitivity and utility of the proposed coding framework using two contrasting dyadic corpora (N=5) of mother-infant object-oriented interactions during experimental conditions that were either conducive (Condition 1) or non-conducive (Condition 2) to the emergence of playful behaviour. We find that the framework accurately identifies the modal form of social interaction as being either playful (Condition 1) or non-playful (Condition 2), and further provides useful insights about differences in the quality of social interaction and temporal synchronicity within the dyad. In conclusion, here, we present a novel neuropsychological framework for analysing the continuous time-evolution of adult-infant play patterns, underpinned by biologically informed state coding along sensorimotor, cognitive and socio-emotional dimensions. We expect that the proposed framework will have wide utility amongst researchers wishing to employ an integrated, multi-level approach to the study of play, and lead towards a greater understanding of the neuroscientific basis of play and may yield insights into a new biologically-grounded taxonomy of play interactions.

## 1 INTRODUCTION

### 1.1 Challenges to a neuropsychological understanding of play

During early life, play is a ubiquitous activity, and engaging in play is positively associated with the development of social skills, cognitive skills, language and emotional well-being (Fung & Cheng, 2017; Lyytinen, Laakso, Poikkeus & Rita, 1999; Pellegrini, Kato, Blatchford & Baines, 2002; St George, Fletcher & Palazzi, 2016; Thibodeau, Gilpin, Brown, & Meyer, 2016). Current conceptualisations of play in the behavioural sciences view it in broad terms as behaviour that is voluntary, engaging, non-functional, and associated with the expression of positive affect (Burghardt, 2005; Lillard et al, 2013; Miller 2017). Play can also be categorised based on the focus of play, i.e. what is the individual playing with? For example, *physical play* is play with one’s own body and other people, for example, climbing, sliding, chasing (Pellegrini et al, 2002; Power, 1999; St George et al, 2016;); *sociodramatic* or *pretend play* is play with a make-believe world, and the focus of play is more than a concrete observable entity (Lillard et al, 2013); *games with rules* involve playing with a set of rules that participants agree to abide by to partake in the play experience, for example, board games or playground games such as tag (Hassinger-Das et al, 2017); and *functional/exploratory play* involves playing with physical objects, such as toys and blocks, and includes constructing models or making things (Pellegrini & Gustafson, 2005; Power, 1999).

These diverse categorisations and definitions show that play in humans is diverse, multi-faceted, and defined by a set of broad terms encompassing motivational, cognitive, social and emotional aspects of behaviour and psychology. Consequently, a challenge for current research is to converge on a common definition and measurement system for play - whether examined at a behavioural, cognitive or neurological level. Central to this challenge is the fact that is difficult to exert the level of experimental control required for investigation at the neurological level, while also retaining the freeform, diverse quality which many consider to be a defining feature of play.

Play behaviour changes substantially across the life-span (Power, 1999), and the model that we present here focuses primarily on play during infancy. Infants primarily engage in the types of play which revolve around physical objects, namely functional and exploratory play, as they do not yet have the cognitive capacities to engage in the more abstract forms such as pretend play or games with rules. As such, the observation of parent– infant object play is a means of studying naturalistic, diverse play behaviour which is empirically viable and theoretically justified. Physical objects occupy an important and developmentally significant position in adult-infant communication. While dyadic interaction - adult-infant - is present throughout the first year of life, it is not until around the end of the first year that infant’s ability to engage in triadic interaction (i.e., adult-infant-object) becomes consolidated (Bakeman & Adamson, 1984; De Barbaro, Johnson & Deak, 2013). This progression to triadic interaction, focussed around an object, is considered important for many aspects of psychological development, including symbolic awareness and language (De Schuymer, Dr Groote, Beyers, Striano, & Roeyers, 2011; Tomasello, 1999). As Rodriguez (2009) points out, objects are symbols of their uses within a culture (a cup, for example, can represent drinking), and an understanding of these object-use relations represents the early acquisition of cultural norms and adoption of a fundamental symbolic system. Adults and infants communicate *about* objects and *with* objects. Consequently, focusing our model around a physical object was deemed the best approach for studying behavioural and neural activity during parent-infant play.

Due to the challenges of experimental control, neuroscience studies on play have primarily focused on animal models (in particular rats) and rough-and-tumble social play behaviour (see reviews by Pellis and Pellis, 2009; Siviy and Panksepp, 2011; Cooke and Shukla, 2011; Vanderschuren et al, 2016). Rodent models have proven to be particularly useful because rats show predictable and stereotypical forms of play-related behaviour (e.g. one animal ‘pins’ the other on its back, emission of ultrasonic vocalisations, etc) which are readily quantifiable and amenable to experimental and pharmacological manipulation. Consequently, a relatively rich literature now exists on the neuroanatomical and neurochemical substrates of rough-and-tumble play behaviour in rats. Namely, the key neural circuits that are now known to work in concert to support rats’ play fighting behaviour are : (1) a ***cortical executive circuit*** (particularly the prefrontal cortex (PFC) and orbitofrontal cortex (OFC)) which mediates the developmental fine-tuning and complexity of play, such as the ability to coordinate with or modify movements in response to the social status of a play partner (Moore, 1985; Pellis et al, 1999; Pellis et al, 2006; Bell et al, 2009; Siviy & Panksepp, 2011); (2) a ***subcortical limbic circuit*** (amygdala, hypothalamus and striatum) which moderates the motivation for, and affective response to play (Burgdorf et al, 2007; Meaney et al, 1981; Daenen et al, 2002; Wolterink et al, 2001), potentially via dopaminergic and opioid pathways (Vanderschuren et al, 2016); and (3) ***somatosensory circuits*** (somatosensory cortex, thalamus, cerebellum) which control motor play patterns and performance (Siviy & Panksepp, 1985, 1987a,b; Panksepp et al, 1994; Byers & Walker, 1995).

Animal studies have further shown that play induces neural plasticity in brain areas involved in sensorimotor processing (e.g. parietal cortex, colliculi and striatum, Gordon et al, 2002), and also in the medial prefrontal cortex (mPFC, Cheng et al, 2008), an area which sends strong modulatory inputs to limbic circuits that control social behaviour. In humans, the mPFC inhibits aggression and monitors approach/avoidance behaviour (Bufkin & Luttrell, 2005; Hall et al, 2010). Therefore, increased plasticity in the mPFC following play could indicate that play helps to improve control of social behaviour networks. Rough-and– tumble play in rats also seems to promote brain development by increasing the expression of brain-derived neurotrophic factor (BDNF) in the amygdala and prefrontal cortex (Gordon et al, 2003), and that of insulin-like growth factor 1 (IGF-1) in the frontal and posterior cortices (Burgdorf et al, 2010). Accordingly, it has been suggested that play-induced neural plasticity could support the emergence of adult-like behaviours (Cooke & Shukla, 2011).

Although caution must be applied in extrapolating findings from animal work to humans, the current data do suggest that across species, play may be a fundamental neurobehavioural process that is underpinned by (and produces changes in) major cortical and subcortical neural circuits that support cognition, emotion and sensorimotor function. However, the neuroscientific methods that have successfully been used with animal models are too invasive to be performed on human subjects. Further, as indicated by the multiple types of play outlined above, human play behaviour is far more complex and less stereotypical than animal play behaviour. Consequently, neuroscience research into play in humans tends to either assess neurological change using a pre-test, post-test design (Newman, Hansen & Gutierrez, 2016), or study neural activity while the participant observes, but does not engage in, play behaviour (Smith, Englander, Lillard & Morris, 2013). Going beyond these empirical constraints to identify the neural mechanisms that underlie ongoing, complex play behaviour in humans presents a considerable challenge. To address this challenge, non-invasive human neuroimaging (e.g. EEG, fMRI) and psychological behavioural coding approaches could be combined into a multi-level analysis that may yield advances in understanding the human neurocognitive mechanisms of play. However, there is currently no integrated framework for conducting a multi-level analysis of play that spans brain, cognition and behaviour. Specifically, there are two major obstacles to achieving the desired coherence across methods.

First, neural analyses require the behaviour of interest to be identified with precise temporal resolution, but existing play coding schemes are predominately based around global ratings, checklists, or frequency counts of play behaviours, and so do not capture temporal information about *when* specific play behaviours occur. For example, The Symbolic Play Test (Lowe & Costello, 1976) consists of a check-list of behaviours which children display when playing with a specific set of toys, such as ‘feeds doll’ and ‘moves truck or trailer about’. A similar checklist approach is found in many bespoke measures of play, such as that used by Pellegrini (1992), where children were observed in the playground and the behaviours displayed were recorded, including *peer interaction* and *object play.* Another common approach is to code various socioemotional dimensions of play globally, i.e., to allocate scores for the entire play session. St George et al (2016) observed parent-child object play and gave each parent a score on 10 different dimensions, including sensitivity (how responsive the parent was to the child’s signals), positive regard (demonstrations of love and affection), and stimulation of cognitive development (teaching). None of these approaches captures specific, temporal information about play behaviour. Therefore, a multi-level coding system for play, suitable for analysing co-occurring patterns in real-time behavioural and neurological data, requires a degree of temporal precision that is not built into existing play coding schemes.

Second (as indicated by animal studies), play is a highly-complex social interactive activity that activates a combination of sensorimotor, cognitive and socio-emotional neural circuits - each of which may support separable dimensions of behaviour. Importantly, no single behavioural dimension by itself is sufficient to define play, since behaviour in each dimension can occur in both playful and non-playful situations. Rather, it is the co-occurrence of activity along multiple dimensions that defines a playful episode. Here, we contribute to the formation of a multi-level neuropsychological understanding of play, through presentation of a model and framework that captures behaviour at a high temporal resolution as a continuously evolving multi-dimensional state, rather than as a set of discrete actions or as a global summary of type or quality. In this way, behavioural coding is well– matched to the high temporal resolution of EEG data, maximising the acuity with which brain-behaviour correlates can be explored.

### 1.2 Overview and considerations of the neuropsychological play coding framework

As described in the previous section, current neuroscientific research suggests that play behaviour is underpinned by three major neural circuits that control motivation and affect (i.e. limbic structures), motor performance (i.e. somatosensory structures), and higher– order executive function (i.e. frontal cortical structures) respectively. Following from this, the proposed coding framework captures object-oriented play behaviour along three corresponding neuropsychological dimensions: the socioemotional dimension (SE), the sensorimotor dimension (SM), and the cognitive dimension (C). Infants’ or adults’ behaviour is coded according to the presence or absence [1/0] of play-related activity in each dimension. Play-related activity in the SE dimension occurs when there is a display of positive or neutral affect, consistent with the idea that play leads to an internal sense of reward (Burghardt, 2005; Miller, 2017). Play-related activity in the SM dimension occurs when the partner (mother or infant) is manipulating and/or touching the object in an exploratory manner, relatively free of external constraint. This criterion reflects the central place of self-directed, voluntary behaviour in definitions of play (Burghardt, 2005; Lillard et al, 2013; Miller, 2017; Sawyer, 2017). Finally, the C dimension captures the presence of attentional engagement, as well as the level of complexity of this cognitive engagement. Therefore, our analysis is intended to explore ‘minds-on’ play, rather than ‘minds-off’ play. According to our framework, a *play-congruent* state is one in which the infant (or adult) concurrently exhibits play-related activity across all 3 dimensions (i.e. [1 1 1]).

An important feature of this framework is that it does not assume any one definition of play. Instead, we have grounded the framework in specific observable behaviours that are considered important factors across different conceptualisations of play - namely, the display of affect and voluntary physical and cognitive engagement with the object of play (Lillard et al, 2013; Miller, 2017). By analysing the co-occurrence patterns of these basic behaviours, our framework can be used to assess similarities and potential groupings of different play– related social states (and, eventually, their neural substrates), which may in future lead to a definition of play behaviour that is grounded in neuroscience. Another strength of the proposed framework is that it does not require the coder to make a subjective judgment about whether or not playful activity is occurring. Rather, objective and observable behaviours are coded (e.g. touching a toy, looking at a toy, smiling, etc), and the presence or absence of play (and other related social states) is inferred from temporally co-occurring patterns of behaviour.

The *joint* social state of adults and infants can also be assessed using this scheme. For example, during didactic teaching, only the cognitive dimension may be concurrently engaged in both partners, whilst their sensorimotor and socio-emotional states may be discordant (e.g. Mother’s state is [1 1 0] whilst the infant’s state is [0 1 1]). Importantly, this framework permits an empirical discrimination between similar/related social interactional states such as teaching versus play. Although the play-congruent state [1 1 1] is of greatest interest here, a total of 8 different social states for infants and adults (and 64 joint adult-infant states) may be discriminated under the proposed framework, which may yield insights into a new biologically-grounded taxonomy of play interactions.

### 1.3 Aims and predictions

Here, we illustrate the application of our proposed neuropsychological play coding framework using examples from two contrasting dyadic corpora of mother-infant object-oriented interactions during experimental conditions that were either conducive (Condition 1) or non-conducive (Condition 2) to eliciting playful behaviour. In Condition 1, playful behaviour was encouraged by asking mothers to use the objects in spontaneous, fun and natural interactions with their child. In Condition 2, playful behaviour was discouraged by asking mothers to focus on teaching infants about the social value (desirable or non-desirable) of the objects. These corpora comprise both behavioural and electroencephalography (EEG) measurements that were collected concurrently from mothersand their infants. However, for this study, we focus solely on behavioural analyses only. We have two specific sets of predictions regarding the differences between conditions that should emerge following application of the coding framework:

1. In Condition 1 (conducive), infants’ modal state will be [1 1 1] (i.e. play-congruent along all three dimensions), but in Condition 2 (non-conducive), [1 1 1] will *not* be the modal state;
2. In Condition 1 relative to Condition 2, infants will show:

a. Decreased negative affect
b. Increased sensorimotor engagement
c. Equivalent cognitive (attentional) engagement

The first prediction pertains to the *sensitivity* of the coding framework in detecting play-related behaviour. Simply put, if mothers were instructed to play with their infants, then (although coders do not make direct judgements about whether participants were playing or not) we expect the coding framework to reveal that a play-congruent state was indeed the most frequent social state that infants displayed. The second set of predictions pertains to the *utility* of the framework in identifying differences in the quality of social interaction and temporal synchronicity with the dyad.

## 2 METHODS

### 2.1 Participants

Five mother-infant dyads participated in the study (3M, 2F infants). Infants were aged 326.6 days on average (range = 292-377 days, SD = 31.5 days). All mothers reported no neurological problems and normal hearing and vision for themselves and their infants.

### 2.2 Materials

For Condition 1, 4 pairs of ambiguous novel objects were used. Within each pair, objects were matched to be globally similar in size and texture, but different in size and colour. Ambiguous novel objects were chosen to ensure that infants would not have their own previous (playful) experience with these objects, and would rely on their mothers’ instruction to guide their interactions with the objects.

For Condition 2, a set of 8 different small toys was used. These were appropriate for the infants’ age and included toys of differing shapes, textures and colours to encourage infants’ interest in playing with them.

### 2.3 Tasks

Each mother-infant dyad took part in experimental Conditions 1 and 2 in a counterbalanced order. In each condition, mothers and infants interacted with objects together, with the major difference being whether the nature of social interaction between mother and infant was conducive to eliciting playful behaviour (as determined by the task instructions provided to the mother). In both tasks, the infant sat in a high chair, with the adult facing him/her across a table. The distance between the infant and adult was the same in each task, and each task lasted approximately ten minutes.

*Condition 1 (not play-conducive)*. In this condition, mothers were asked to teach their infants about the social value of pairs of ambiguous novel objects. For each pair of objects, mothers were instructed to describe one object with positive affect (“This is great, we really like this one!”) and the other object with negative affect (“This is bad, we don’t like this one”), as shown in Figure 1. Mothers were asked to limit their verbal descriptions to four simple formulaic sentences per object (which they repeated for each object), and to model positive or negative emotions in a prescribed manner (e.g. smiling versus frowning). The order of object presentation (positive or negative) was counterbalanced across trials. After observing their mothers’ teaching about both objects, infants were then allowed to interact briefly with the objects themselves before the objects were retrieved. During the session, an experimenter was present to ensure that participants were interacting as instructed. She provided new pairs of objects as required, but explicitly avoided making prolonged social contact with either participant.

**Figure 1.**
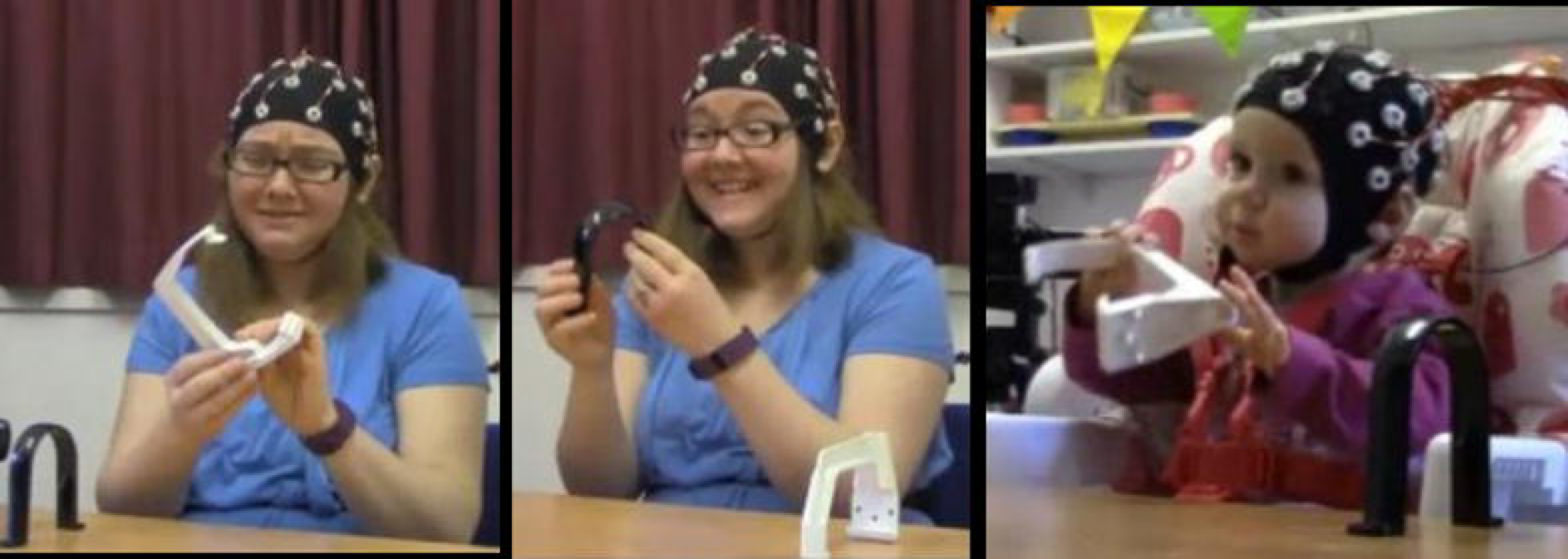
*Illustration of experimental setup for Condition 1*. *(left) Negative object demonstration by adult; (middle) positive object demonstration by adult; (right) infants’ interaction with objects*.

*Condition 2 (play-conducive)*. In this condition, mothers were asked to play with their infant using a set of attractive toys (see Figure 2). Mothers were instructed to use the toy objects in a spontaneous, fun and natural way, to actively engage the infant’s attention, but to play quietly whilst avoiding large physical motions (in order to minimise EEG motion artifacts). During the session, an experimenter was present to ensure that participants were playing as instructed. She provided new toys as required (approximately every two minutes, or more frequently if the child threw the object to the floor) to sustain their attention and interest. The experimenter explicitly avoided making prolonged social contact with either participant.

**Figure 2.**
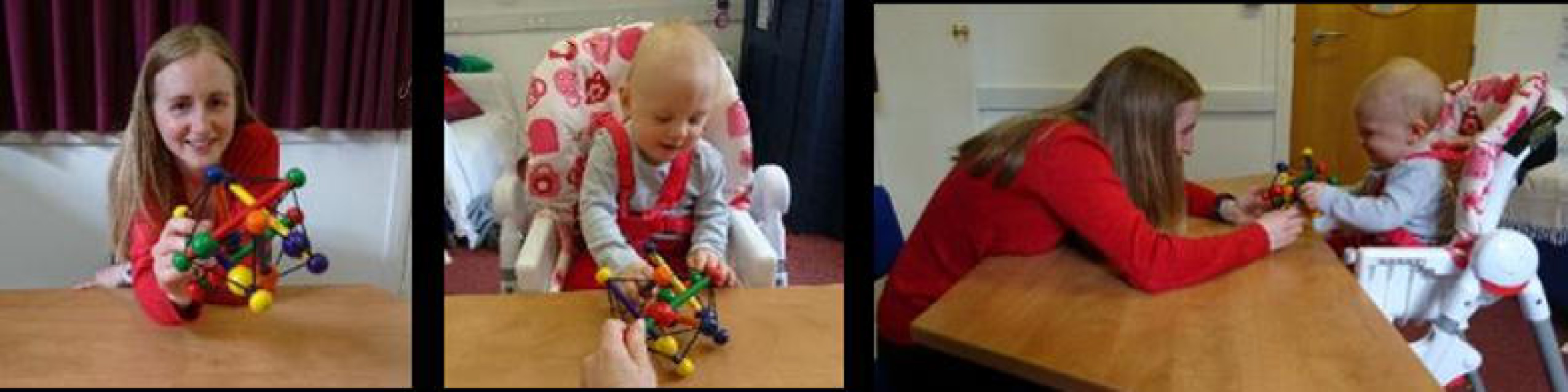
*Illustration of experimental setup for Condition 2*. *(left) Infants’ view; (middle) adults’ view; (right) side view of social interaction. Note that although the actors here are not wearing EEG caps, EEG signals were also collected during this condition*.

### 2.4 Video recordings

To record the actions of the participants, two Logitech High Definition Professional Web-cameras (30 frames per second) were used, directed at the adult and infant respectively. Afterwards, each video recording was manually coded for the timing of the behaviours of interest, using the coding scheme outlined in Section 2.5.

EEG data was also concurrently collected from mothers and infants during social interactions, but this data is not reported here as the primary focus of the current study is to develop a framework for assessing play behaviour.

### 5 A neuropsychological framework for analysing adult-infant play patterns

#### *2.5.1* *Coding scheme*

The coding scheme was designed to capture object-oriented play behaviour in three neuropsychological dimensions: the socioemotional dimension (SE), the cognitive dimension (C), and the sensorimotor dimension (SM). On each dimension, simple, observable behaviour at each timepoint (here, at the temporal resolution of 33ms, corresponding to 30 frames per second) is coded using a [1/0] main code which indicates the presence or absence of play-congruent activity along the target dimension. Additionally, and where relevant, a further sub-code [1/0.x] may be assigned to indicate the level/type of activity that is occurring. The coding scheme is summarised in Table 1 below.

**Table 1.**
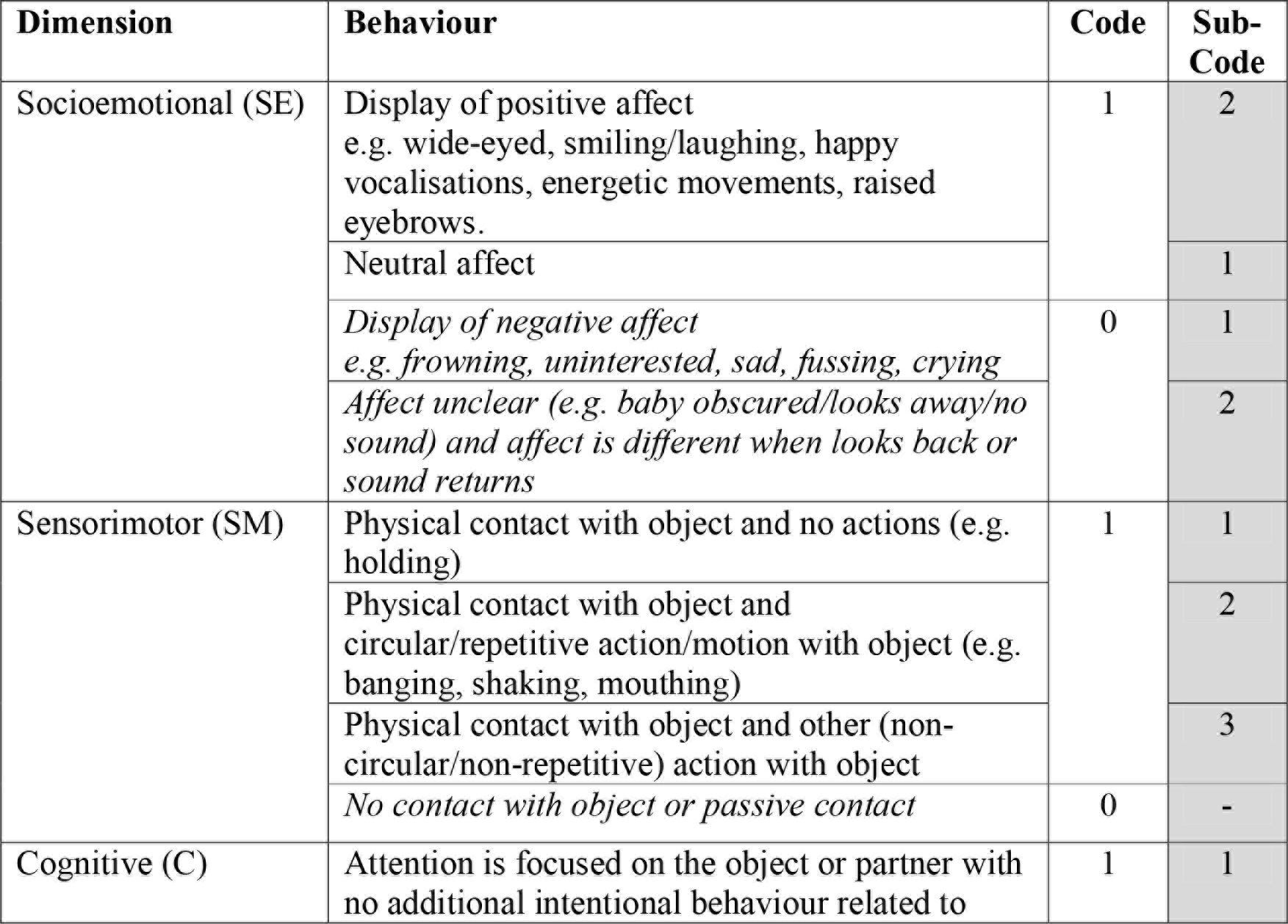
*Coding scheme*

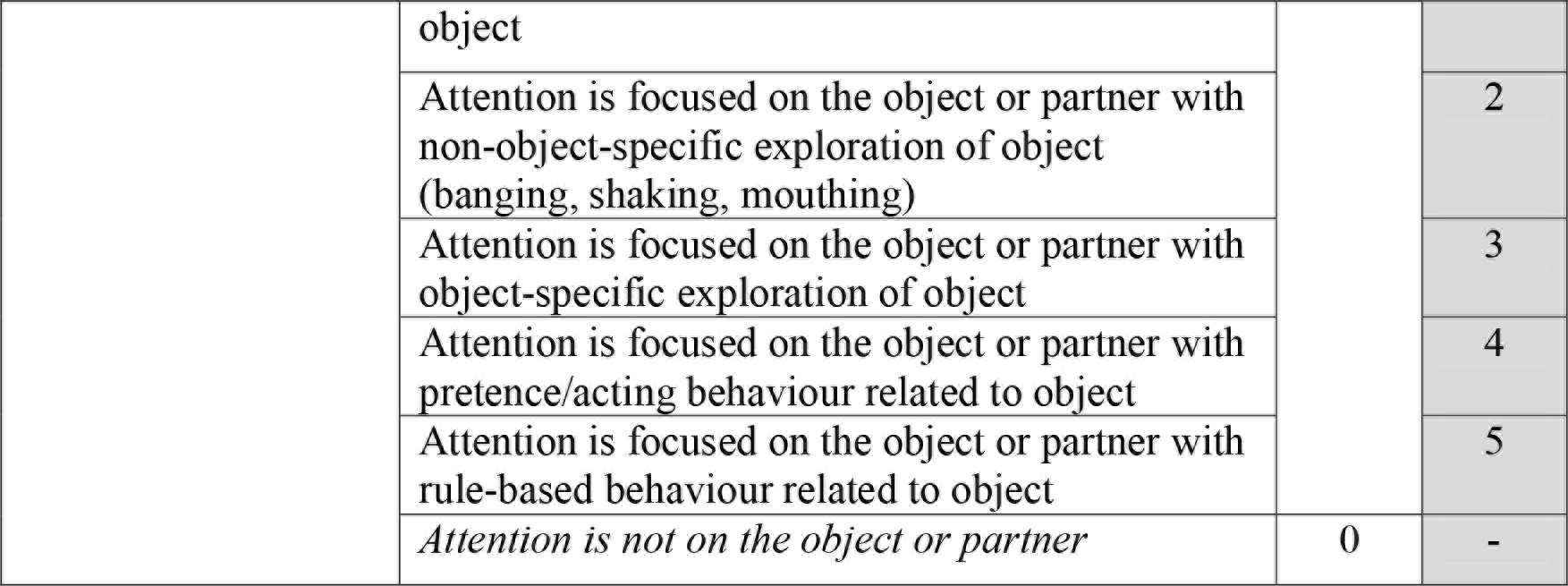

The term ‘play-congruent’ is used to refer to behaviours and states in each dimension during which play might be occurring, and where the individual might be in a playful mental frame. Initially, we sought to base our model around the broadest criteria for a play-congruent state, and then ensure that states based on narrower criteria could also be discriminated. In each dimension, the presence of play-congruent behaviour is allocated a code of 1 and the absence of play-congruent behaviour is allocated a code of 0 (see ‘Code’ column in Table 1). When play-congruent behaviour is *concurrently* observed across all three dimensions (i.e. [1 1 1]), the resulting state is termed a ‘play-congruent state’. This model can be used to explore how individuals move into and out of these broadly-defined play-congruent states (and in future, to explore related neural activity), as illustrated in Section 2.6.

Note that due to the flexibility of the model, it is also possible to analyse how individuals move into and out of narrower types of social states, for example, one could restrict the analysis to only those states involving the expression of positive affect, a high level of cognitive engagement, and physical activity with the object. Our intention in developing the model was to provide researchers with a tool to compare many different interactional states at a behavioural and neural level, with the ultimate goal of developing a neuropsychological taxonomy of play.

*Socioemotional (SE)*. The presence of positive affect and the idea that play is done for its own sake, leading to an internal sense of reward rather than any form of external reward, are both central to most conceptualisations of play behaviour (Burghardt, 2005; Miller, 2017). However, whilst negative affect is considered antithetical to the presence of a mental ‘play-state’, a neutral display of affect could also be present during play (Miller, 2017). Therefore, the expression of positive or neutral affect was taken as congruent with a play-state in the socioemotional dimension.

*Sensorimotor (SM)*. The sensorimotor dimension captures whether or not there is voluntary physical contact with the object that is free from external constraint. It is not possible to engage in object play without physical contact with the object, so this dimension encodes a necessary condition for one of the main play behaviours of interest during infancy. Furthermore, the fact that only voluntary contact is coded reflects the central place of self-directed, voluntary behaviour in definitions of play (Burghardt, 2005; Lillard et al, 2013; Miller, 2017; Sawyer, 2017). Therefore, voluntary physical contact with the object was deemed as congruent with a play-state in the sensorimotor dimension.

*Cognitive (C)*. The cognitive dimension captures the level of cognitive complexity and engagement, by coding whether or not there is visual attention on the object and/or play partner and what kind of behaviour is occurring in relation to the object. While playful behaviour may occur without active attention on the object or play partner (for example, an infant swinging a toy around while not looking at anything in particular, or with eyes closed), we decided that, as we were interested in play’s effects at the neural level, even our broadest criteria for a play-congruent state should include some level of cognitive engagement. In other words, our model is intended to explore ‘minds-on’ play, rather than ‘minds-off’ play that is purely physical or sensory in nature. Therefore, visual attention on either the object or the play partner was deemed as congruent with a play-state in the cognitive dimension.

In the cognitive dimension five sub-codes were also developed to delineate whether the individual is also engaged in exploratory behaviour (object-general or object-specific), pretence or acting, or rule-based behaviour. The distinction between object-general and object-specific exploration is intended to capture two different levels of cognitive engagement which are expected to relate to observable differences in neural activity. Object-general exploration is any kind of activity with the object that does not involve appreciation of the object’s particular properties, i.e. the action could be done with almost any object. The main examples of object-general exploration include shaking, banging, or mouthing the object, and these behaviours are often done in a repetitive or circular fashion. Object-general exploration may provide sensory stimulation and coarse, ‘global’ information, but seems unlikely to lead to specific conceptual information about an object’s functions and uses. Object-specific exploration, by contrast, involves an appreciation of that object’s unique properties - for example, spinning the blades of a toy helicopter, pushing on the surface of a balloon, or pulling on parts of the object to see if they can be removed. It is this kind of exploration that seems most likely to lead to more advanced conceptual learning about an object’s functions and uses. Figure 3 shows an example of the resulting codes for each separate dimension over time during a social interaction episode for an infant:

**Figure 3.**
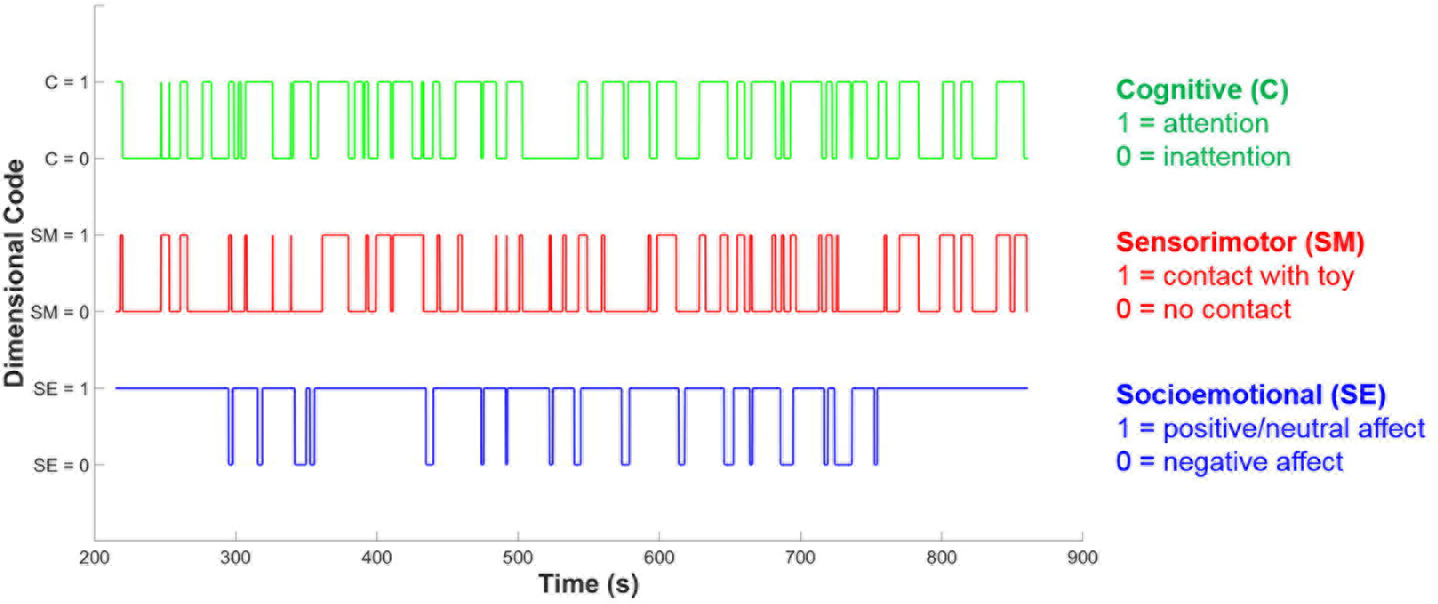
*Example of neuropsychological dimensional coding over time*. *The blue line shows socioemotional (SE) dimension coding, the red line shows sensorimotor (SM) dimension coding and the green line shows cognitive (C) dimension coding*. *At each time point, each dimension may either be coded as 0 or 1, indicating the absence or presence of play– congruent dimensional activity*.

### 2.6 Analysis of social states

#### *2.6.1* *Mean dimensional scores*

For each dimension (socioemotional [SE], sensorimotor [SM], cognitive [C]), separate SE, SM and C mean dimensional scores can be computed by taking the average over all timepoints in the session. This mean score ranged between 0 (if all timepoints were coded as 0) and 1 (if all timepoints were coded as 1). Accordingly, if infants generally displayed more positive/neutral affect than negative affect, their SE mean score would be greater than 0.5 (i.e. higher proportion of 1s than 0s overall). Similarly, the mean SM score indicates the proportion of time during which the infant has active “hands-on” possession of the toy (e.g. mean SM score of 0.7 = infant has active possession of the toy 70% of the time). Finally, the mean C score indicates the relative attentiveness of the infant during the session (e.g mean C score of 0.6 = infant is attentive toward the object or partner 60% of the time). It is important to emphasise that a high score on a single dimension (or indeed across several dimensions) does not in itself indicate that the infant is highly engaged in play, since these mean dimensional scores are independent of each other and therefore provide no information about *temporal co-occurrence* across dimensions (i.e. social state). For example, it would, in theory be possible for the infant to show positive affect *only when* inattentive/not touching the toy and yet still produce a high mean score on the SE dimension.

#### *2.6.2* *Social states (per timepoint)*

At each timepoint, an infant’s current social state can be defined by the temporal co-occurrence and valence of existing codes on each dimension, signifying affect (SE), touch (SM) and attention (C) respectively. As SE, SM and C codes could each take a value of 0 or 1, this allows a total of 8 distinct social states (i.e. 2^3^), as outlined in Table 2 and Figure 4.

**Table 2.** *Examples of possible social states and their potential interpretation*

| Social State | Dimension Main Code |  |  | Interpretation |
| --- | --- | --- | --- | --- |
|  | SE | SM | C |  |
| 1 | 1 | 1 | 1 | Positive/neutral affect, contact and attention<br>(= ***Play-congruent state***) |
| 2 | 1 | 0 | 1 | Positive/neutral affect, no contact, attention |
| 3 | 0 | 1 | 1 | Negative affect, contact, attention |
| 4 | 0 | 0 | 1 | Negative affect, no contact, attention |
| 5 | 1 | 1 | 0 | Positive/neutral affect, contact, inattention |
| 6 | 1 | 0 | 0 | Positive/neutral affect, no contact, inattention |
| 7 | 0 | 1 | 0 | Negative affect, contact, inattention |
| 8 | 0 | 0 | 0 | Negative affect, no contact, inattention |

**Figure 4.**
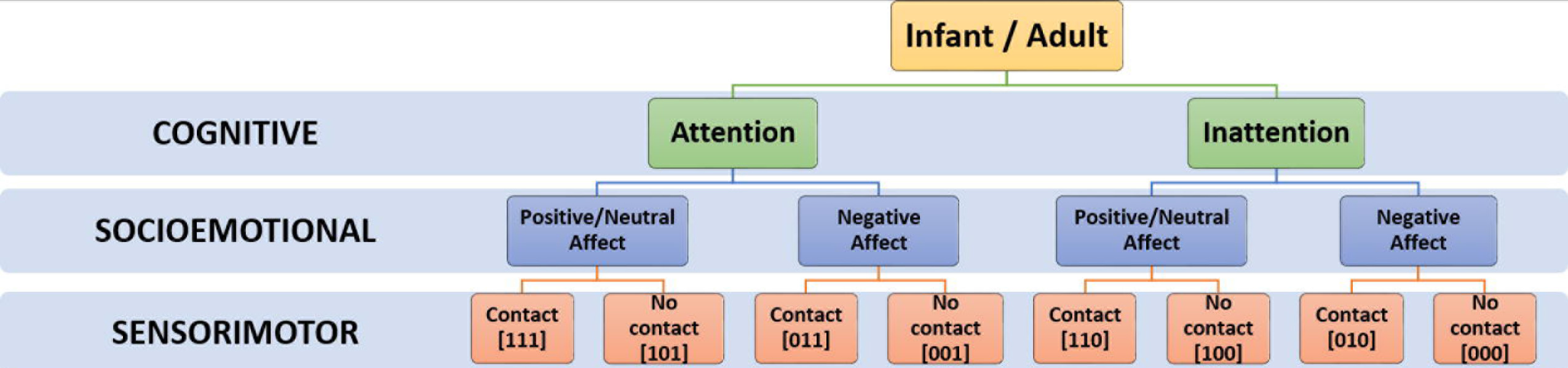
*Hierarchical taxonomy of possible social states*

Further, as social states were computed for every time-point in the session, this permits the tracking of infants’ dynamic evolution between social states over time, as illustrated in Figure 5a, as well as the relative proportion of time that infants spend in each state (Figure 5b. Finally, using the state frequency histogram, it is possible to identify the ***modal*** social state for a given interaction session.

**Figure 5.**
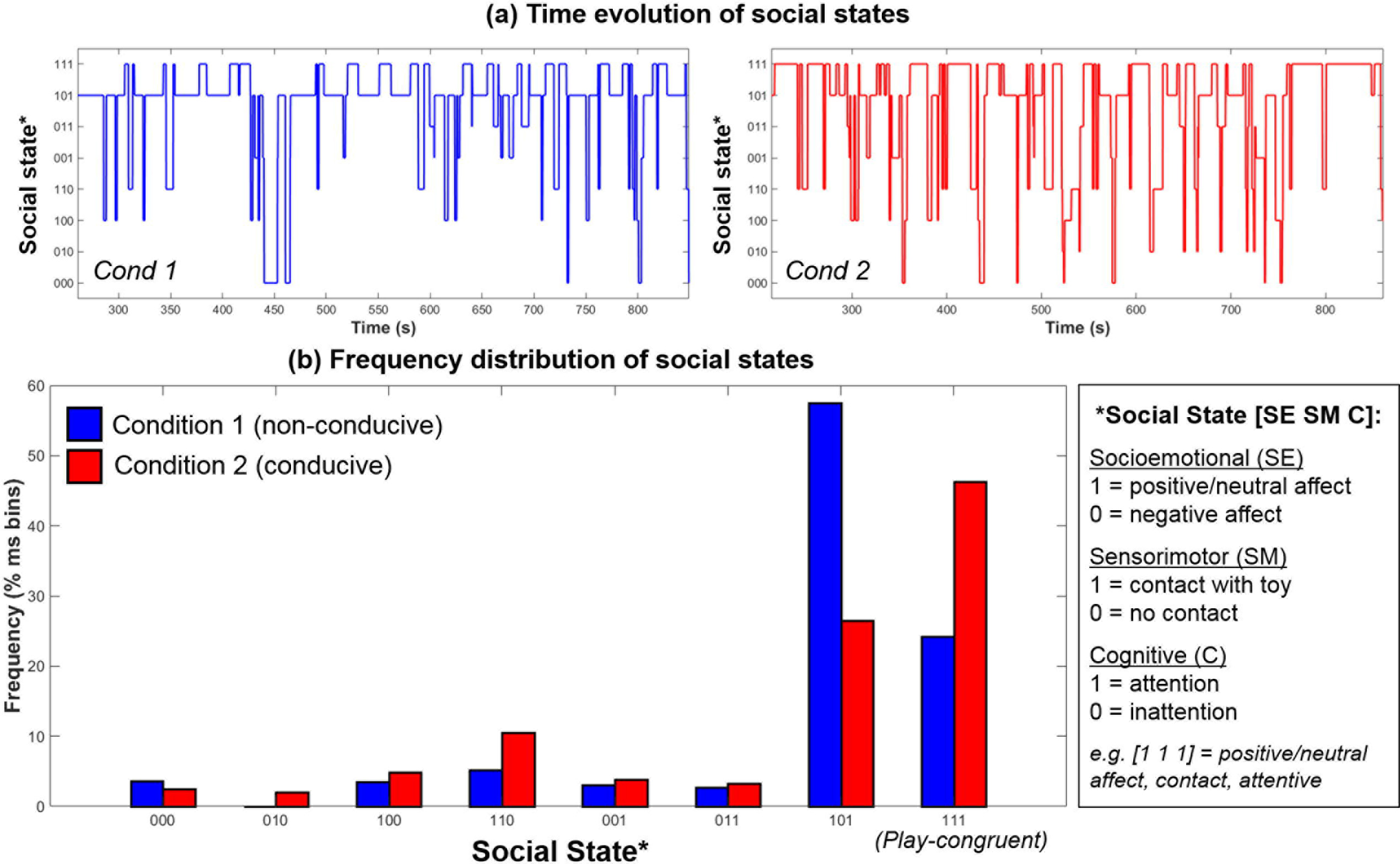
*Example data from one infant showing the (a) Time-evolution between 8possible social states during Condition 1 (left, blue) and Condition 2 (right, red); (b) Frequency distribution of social states in Condition 1 (blue) and Condition 2 (red)*

#### *2.6.3* *Mother-infant joint states (behavioural state synchrony)*

As a further step, both mother’s and infant’s behaviour may be coded using this scheme and their *joint* (i.e. concurrent) social states may be identified. Again, this may be performed to examine joint states within a particular dimension (e.g. affect), considered alone. Or up to 64 (8 × 8) different joint states may be discriminated, which could be useful to address research questions pertaining to parent-child synchrony (since if both parent and child display the same state at the same time, they are behaving synchronously), contingency and responsiveness.

## 3 RESULTS

Here, we report the results from the *main* codes assigned along each dimension (e.g. 1 or 0 - see Table 1). However, if desired, more fine-grained information about the quality of social interaction may be gleaned by examining participants’ sub-codes, as detailed in the Supplementary Materials.

### 3.1 Mean dimensional scores

Infants’ and mothers’ mean dimensional scores obtained during Conditions 1 and 2 are shown in Figure 6. For each group of participants (infants and adults), differences between conditions were assessed statistically using a Repeated Measures ANOVA taking Dimension (3 levels, SE/SM/C) and Condition (2 levels) as the within-subjects factors.

*Infants*. The ANOVA revealed that there was a significant main effect of Condition (F(1,4) = 17.65, p<.05, η^2^p = .82), where, across all dimensions, mean dimensional scores for Condition 2 (play-conducive) exceeded those for Condition 1 (not play-conducive). Importantly, there was also a significant interaction between Condition and Dimension, suggesting that not every dimension differed between Conditions (F(2,8) = 8.80, p<.01, η^2^p = .69). Tukey HSD *post hoc* analysis of this interaction indicated that Socioemotional and Sensorimotor dimensional scores were both significantly higher during Condition 2 than Condition 1 (p<.05, p<.01 respectively), but Cognitive dimensional scores did not differ (p = .99). Therefore, during social interactions that supported playful behaviour, infants (on average) showed more positive/neutral affect and active possession of the toy, but they were equally attentive across both conditions.

**Figure 6.**
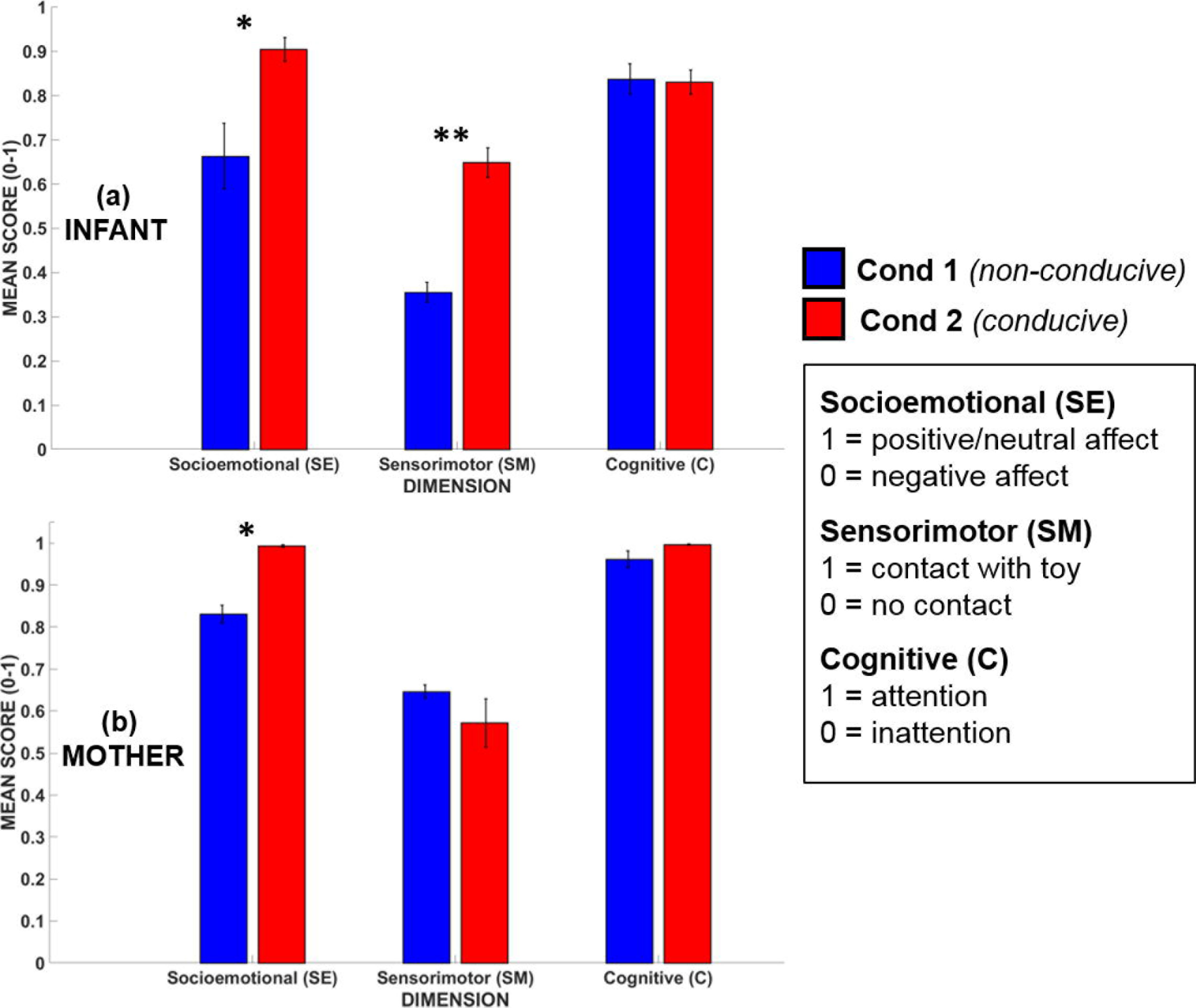
*Mean dimensional scores for (a) infants’ and (b) mothers’ in Condition 1 (blue) and Condition 2 (red)*. *Error bars indicate the standard error of the mean*. *p<.05, **p<.01

*Mothers*. The ANOVA revealed no significant main effect of Condition (F(1,4) = 2.24, p=.21, η^2^p = .36). However, there was again a significant interaction between Condition and Dimension, suggesting that not every dimension differed between conditions (F(2,8) = 11.1, p<.01, η^2^p = .74). Tukey HSD post hoc analysis of this interaction indicated that, like infants, mothers also showed significantly greater expression of positive/neutral affect (i.e. scores on the Socioemotional dimension were higher) during Condition 2 as compared to Condition 1, but mothers’ mean scores on the other two dimensions did not differ between conditions.

Whilst dimensional scores are able to capture time-averaged differences in overall social interactional quality, they cannot reveal whether qualitatively-different *types* of social interaction are occurring (as well as their timing and frequency of occurrence). Accordingly, we next assessed infants’ social states (calculated for each timepoint) in each condition.

### 3.2 Frequency distribution of social states during play and teaching

The frequency distribution of different social states observed in infants and mothers during Conditions 1 and 2 are shown in Figure 7. Given that there were only 5 data points, this provided insufficient degrees of freedom to conduct an omnibus Repeated Measures ANOVA. Accordingly, to assess whether there were statistical differences between conditions in social state frequency, we conducted paired t-tests (Benjamini-Hochberg FDR corrected p-values, alpha = .05) for each social state, for infants and adults.

*Infants*. The t-test results revealed that there was a large and significant increase in the frequency of the play-congruent [1 1 1] social state during Condition 2 as compared to Condition 1 (t(4) = 8.97, BH-FDR p<.01). On average, during Condition 1, infants were in a [1 1 1] state 24.9% of the time but during Condition 2, this frequency doubled to 50.7% (i.e. half) of the time spent in social interaction. Similarly, during Condition 2, there was a marginal (near-significant) increase in the frequency of the [1 1 0] social state (t(4) = 2.90, BH-FDR p =.12), and marginal decrease in the frequency of the [0 0 1] social state (t(4) = − 3.11, BH-FDR p =.12). Together, these results suggest that during Condition 2 (which was conducive to playful behaviour), infants spent significantly more time in social states characterised by “positive-affect hands-on interaction” (i.e. [1 1 1] or [1 1 0]), and proportionately less time in a negative-affect passive observational state [0 0 1].

**Figure 7.**
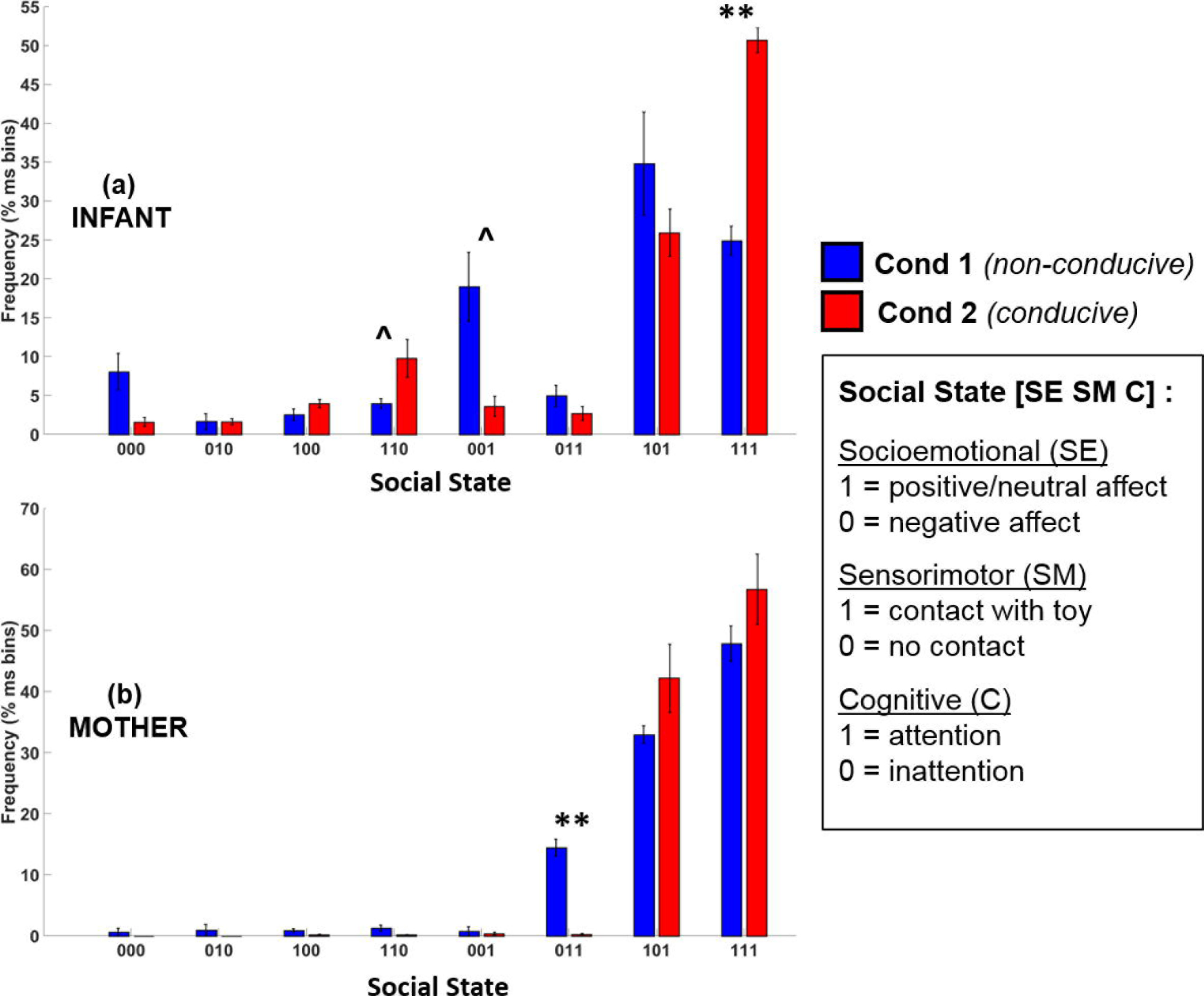
*Frequency distribution of the eight possible social states for (a) infants and (b) mothers across Condition 1 (blue) and Condition 2 (red)*. *Error bars indicate the standard error of the mean. **p<.01, ˄p=.12 (BH-FDR corrected)*

*Mothers*. Mothers’ t-test results revealed only one significant difference between conditions. There was a significant decrease in the frequency of the play-incongruent socialstate of [0 1 1] (negative affect, contact, attention) in Condition 2 as compared to Condition 1 (t(4) = -9.85, BH-FDR p<.01). However, although there was a trend toward an increase in play-congruent behaviour ([1 1 1]) in mothers for Condition 2, this increase was not significant (t(4) = 1.28, BH-FDR p=.43). Therefore, although mothers displayed less play-<u>*in*</u>congruent behaviour during Condition 2 than Condition 1, we did not observe significantly more play-congruent behaviour.

### 3.3 Individual modal states

*Infants*. During Condition 1, the modal (most frequently-occurring) social state was [1 0 1] for 4 infants, and [0 0 1] for 1 infant. However, during Condition 2, the modal social state for all 5 infants was the play-congruent state of [1 1 1]. Therefore, there was a clear difference in the characteristic social state of infants between conditions. During social interactions that were non-conducive to playful behaviour, infants were predominantly passive (“hands-off”) but attentive. During social interactions that supported playful behaviour, infants were predominantly active (“hands-on”), positive and attentive.

*Adults*. By contrast, mothers displayed almost no difference in their modal states across experimental conditions. Four out of five mothers showed a modal state of [1 1 1] for both Conditions 1 and 2. One mother showed a modal state of [1 1 1] during Condition 1, and a modal state of [1 0 1] during Condition 2, with [1 1 1] being her next most frequently– occurring social state.

### 3.4 Mother-infant joint states (behavioural state synchrony)

Finally, we assessed the joint probability distribution of infants’ and mothers’ social states during Conditions 1 and 2, as shown in Figure 8. Of note, perhaps the most relevant difference is that during Condition 1, mothers and infants were in a joint play-congruent social state (i.e. [1 1 1] - [1 1 1], or *synchronous play)* only 5.7% of the time on average. By contrast, during Condition 2, mothers and infants showed synchronous play 24.9% of the time — a nearly five-fold increase. Therefore, during conducive social contexts (Condition 2), the play-congruent state occurred more frequently in regard to infants’ own behaviour, and this joint social state also occurred concurrently (i.e. synchronously) with their mothers more often.

**Figure 8.**
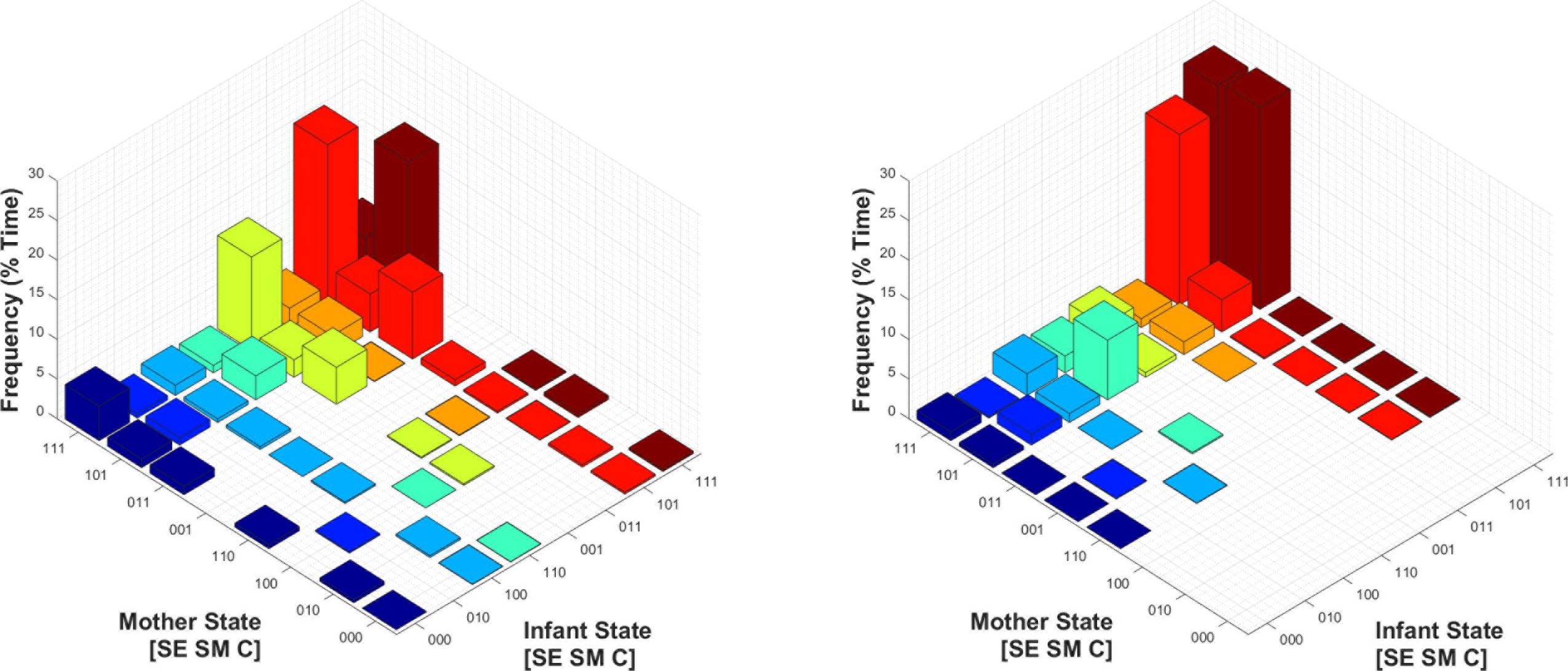
*Mean joint probability distribution of mothers’ and infants’ social states during Condition 1 (left) and Condition 2 (right)*. *The joint play-congruent state of [1 1 1] - [1 1 1] for mother and infant respectively is shown in the posterior corner of each subplot, and shaded in dark red*.

## 4 DISCUSSION & CONCLUSION

Play behaviour is diverse and multi-faceted, and a major challenge for current research is to converge on a common definition and measurement system for play that integrates behavioural, cognitive and neurological levels of analyses. Here we present and test a new methodological framework that captures different social interactional states (play-congruent or play-incongruent) and permits an empirical discrimination between similar and related social interactional states (such as joint activities in situations both conducive and non-conducive to the emergence of playful behaviour).

A-priori, we made two sets of predictions about the differences in infants’ behaviour between these conditions and our coding results supported both predictions. First, we observed that during conducive social interactions, infants’ modal state was indeed play-congruent (i.e. [1 1 1]). Further, infants spent significantly more time in social states characterised by “positive-affect hands-on interaction”, and proportionately less time in a negative-affect passive observational state. This result demonstrates the *sensitivity* of the coding scheme in correctly identifying the intended mode of social interaction as either playful or non-playful, even though coders did not explicitly code for play itself. Further, our data also highlight the fact that, although mothers were instructed to play with their infants during Condition 1, infants themselves did not display playful behaviour *all of the time* (on average, only 50.7% of the time). Rather, infants showed a heterogenous mixture of social states characterised variously by positive affect and “hands-on” engagement. This behavioural finding has important practical implications for the analysis of infants’ *neural* data that is collected during such play sessions. If infants’ neural data during play is assumed to be homogenous and analysed as such (e.g. by computing averages of neural indices across all time-points), such an analysis would be erroneous as it would include many periods when play is not actually occurring. The proposed coding framework therefore lends itself well to time-sensitive neural analyses, because it permits the automatic extraction of discrete time periods when a playful social state is observed (i.e. [1 1 1]), as well as separate analyses of the periods leading up to, and away from these moments.

As predicted, our coding results also showed that, when infants were engaged in social interactions that were conducive for playful behaviour (Condition 2), they showed decreased negative affect, increased sensorimotor involvement with objects, but equivalent attentional engagement. Similarly, mothers also showed significantly decreased negative affect during social interactions that supported playful behaviour. This result demonstrates the potential utility of the coding scheme in highlighting key differences between different forms of early social interactions. Specifically, the finding that infants were no less cognitively engaged during interactions where the mother was not explicitly teaching her infant suggests that playful interactions might provide an equally (if not more) effective social context for early learning as compared to direct didactic instruction from parents.

Finally, we observed that during play, parent-child dyads showed greater temporal synchrony with each other’s social states, as mothers and infants were concurrently (jointly) in a playful state (i.e. [1 1 1] - [1 1 1]) five times more frequently than was observed during teaching. This strong alignment of social-affective state between parent and child during playful scenarios is consistent with previous work. For example, patterns of temporally synchronous activity between parent and child during social interaction have been noted for gaze (Kaye & Fogel, 1980), affect (Cohn & Tronick, 1988; Feldman et al, 2011) and even autonomic arousal (Feldman et al, 2011; Waters et al, 2014). Our coding scheme not only allows the identification of specific time periods when play is synchronously occurring between mother and child (e.g. for neural analyses), but also allows comparison to periods when the dyad is socially asynchronous, or ‘out of tune’ with each other. Such parent-child asynchrony is known to occur more frequently and to be of particular clinical relevance in affective disorders such as maternal depression (Goldsmith & Rogoff, 1997; Jameson et al, 1997).

One major limitation to the current study is its small sample size. However, our intention has been to illustrate the sensitivity of the proposed framework in discriminating between different play-related states, using a coding scheme grounded in simple, observable behaviour and with a temporal resolution suited to neuroscientific research. Five dyads provided sufficient data to assess the efficacy of the framework as a means of capturing variations in individual and joint behaviour as continuously evolving states, rather than as discrete actions or a subjective global assessment. Nevertheless, research applying our framework to larger samples is needed to ensure that the contextual differences we identified between play and teaching scenarios are generalisable.

A second limitation is our focus on one specific type of play which revolves around a physical object. However, the focus on a physical object does not limit our model entirely to the earlier and more basic types of play, because a physical object can be used in play with more symbolic content. For example, object substitution ‒ using an object as if it is something else ‒ is often coded in established play coding schemes as an indicator of pretend play. Many games with rules involve physical objects, so participants could engage in such a game, either spontaneously or because they are asked to do so. We decided to capture these more complex types of behaviour in our model, with the acknowledgement that in infancy, these types of behaviour will be very rare, and most likely observed on the part of the parent. Also, with appropriate development (e.g. the elaboration of sub-code options) our framework could be applied to more abstract forms of play that do not revolve around physical objects.

A final limitation is that, as play behaviour changes significantly across the life-span (Power, 1999), we chose to focus primarily on infants. Therefore, adult play behaviour may not have been optimally-captured by the framework. Nonetheless, our coding revealed an interesting result: mothers (unlike their infants) did not show a clear shift toward greater playfulness for Condition 2 as compared to Condition 1, although their negative affect decreased overall. One possible reason for this may be that mothers approached the teaching exercise in Condition 1 as pretend play. Since the objects used in Condition 1 had no intrinsic social value, mothers had to act out a ‘good’ or ‘bad’ response to the objects, by pretending that the objects had a particular social significance. However, it is clear from our data that infants (aged on average 10.7 months) responded less playfully to their mother’s social pretend play, consistent with the late emergence of pretend play capabilities during the second year of life (Fein, 1981).

Although the neural EEG data that was collected during the parent-child social interactions was not analysed here, the proposed framework represents an important first step towards conducting meaningful analyses of these neural data. Specifically, since the proposed framework fractionates play behavior along neuropsychological dimensions, this provides the capability for (1) internal prediction and validation (e.g. during periods of sensorimotor activity only, neural activity should predominantly arise from the sensorimotor cortices) and (2) the identification of common neural substrates that underlie play-related social interactional states. An analysis of these underlying neural substrates may, in future, lead to a definition of play behaviour that is more closely grounded in neuroscience. The further subcategories of different types of cognitive interaction also included in the scheme (see Table 1) will also allow us to look, in more detail, at other types of play. For example, it may be of interest to assess whether the use of *pretend play* by parents (for example, when the mother pretends that a ball is an animal) is related to the early development of pretense capabilities in infants. In adults, the observation of substitute object pretense leads to activity in the superior parietal lobule (Smith et al, 2013). If the observation of object substitution in infancy was also found to lead to similar patterns of neural activation, this may provide neurological evidence for the facilitation of an emerging awareness of pretence in infants in response to parental pretend play.

In conclusion, we have presented a novel neuropsychological framework for analysing the continuous time-evolution of adult-infant play patterns, underpinned by biologically informed state coding along sensorimotor, cognitive and socio-emotional dimensions. We expect that the proposed framework will have wide utility amongst researchers wishing to employ an integrated, multi-level approach to the study of play, and lead towards a greater understanding of the neuropsychological basis of play.

## ACKNOWLEDGEMENTS

This research was funded by an ESRC Transforming Social Sciences collaboration grant (ES/N006461/1) to VL and SW, and by ESRC FRL Fellowship (ES/N017560/1) to SW.

